# Bivalent CK2 Inhibition as a Potent Strategy Against SARS-CoV-2 and Other Coronaviruses

**DOI:** 10.64898/2026.10.05.756735

**Authors:** Dennis Salomon Lopez Molina, Nazim Ahnou, Rozenn Brillet, Jean-Guy Delcros, Laurent Softic, Bryan Jimenez Araya, Maxime Duval, Patrice Bruscella, Mario Veruete, Alexandre Bancet, Dylan Grenier, Angelique Mularoni, Jean-Michel Pawlotsky, Isabelle Krimm, Abdelhakim Ahmed-Belkacem

## Abstract

Protein phosphorylation is a critical host-signaling pathway hijacked by viruses to facilitate their replication. Casein Kinase 2 (CK2), a ubiquitously expressed serine/threonine kinase, has emerged as a key host factor exploited by several viruses, including SARS-CoV-2. Here, we investigate the role of CK2 subunits (CK2α, CK2α’, and CK2β) in SARS-CoV-2 replication and evaluate the antiviral potential and broad-spectrum anti-coronavirus activity of two CK2 inhibitors: the previously described ATP-competitive inhibitor CX4945 and a newly developed, highly selective bivalent inhibitor, AB668.

SiRNA-mediated knockdown of CK2α, CK2α’, and CK2β significantly reduced SARS-CoV-2 RNA production, nucleoprotein expression, and viral titers, highlighting the essential role of CK2 in viral replication. Co-immunoprecipitation assays revealed that the SARS-CoV-2 nucleoprotein preferentially interacted with CK2α and CK2β rather than CK2α’. Furthermore, bio-layer interferometry assays demonstrated that the nucleoprotein specifically binds to the tetrameric CK2 holoenzyme (CK2α2β2) but not to individual CK2α or CK2β subunits.

Pharmacological inhibition of CK2 with AB668 exhibited potent antiviral activity, significantly reducing viral replication. Time-of-addition experiments indicated that AB668 targets a pre-replicative or RNA-replicative step of the viral life cycle. Importantly, AB668 displayed broad-spectrum antiviral activity, effectively inhibiting both HCoV-OC43 (β-CoV) and HCoV-229E (α-CoV).

In summary, our findings establish CK2 as a critical host factor for SARS-CoV-2 replication and identify AB668 as a promising candidate for broad-spectrum antiviral therapy. With its high selectivity, minimized off-target effects, and efficacy against multiple coronaviruses, AB668 represents a valuable chemobiological and therapeutic tool.

## INTRODUCTION

Protein phosphorylation is a ubiquitous host-signaling pathway that viruses hijack to facilitate their replication. During viral infection, cellular kinases are often activated, leading to the phosphorylation of both viral and host proteins. Casein Kinase 2 (CK2), one of the most pleiotropic kinases, contributes to approximately 24% of the phosphoproteome [1]. CK2 is a ubiquitously expressed, constitutively active serine/threonine kinase that localizes primarily in the nucleus and cytoplasm, where it regulates essential cellular processes such as signal transduction, cell cycle progression, and apoptosis [2, 3].

CK2 functions as a tetrameric holoenzyme composed of two catalytic subunits (α or α’), encoded by the CSNK2A1 and CSNK2A2 genes, respectively, and two regulatory β subunits, encoded by the CSNK2B gene [2]. Different holoenzyme combinations exist, including α₂β₂, α’₂β₂, and αα’β₂. While the catalytic activity of CK2α subunits is independent of CK2β, the regulatory subunit significantly influences CK2 substrate specificity, localization, and stability [4]. Although CK2α and CK2α’ share overlapping functions, they also exhibit distinct roles [5] [4]. These differences primarily arise from variations in their C-terminal sequences, which allow them to bind different partner proteins, thereby conferring isoform-specific functions [5, 6 32].

CK2 has been implicated in the phosphorylation of a wide range of viruses including papillomavirus E7 [7], HIV-1 [8], hepatitis C [9] and B [10], Rotavirus [11], Respiratory syncytial virus (RSV) [12], Herpes Simplex Viruses (HSV) [13] and Human Cytomegalovirus (CMV) [14]. Recent mass spectrometry-based proteomic and phosphoproteomic studies have identified interactions between the nucleoprotein of SARS-CoV-2 and host CK2 in SARS-CoV-2 infected cells [15, 16]. SARS-CoV-2 infection has been shown to activate CK2 activity and increase phosphorylation levels of CK2-substrates, suggesting a functional role of CK2 in SARS-CoV-2 life cycle. Furthermore, these observations extend to

SARS-CoV and MERS-CoV, suggesting that CK2-nucleoprotein interactions are likely conserved among β-coronaviruses [17]. Additionally, Yang et *al.* employed a chemogenomic approach using multiple ATP-competitive CK2 inhibitors to demonstrate that CK2 inhibition suppresses the replication of murine, bat, and human β-CoV [18]. These findings highlight the potential of targeting CK2 as a broad-spectrum antiviral strategy against β-coronaviruses.

ATP-competitive inhibitors targeting CK2α’s catalytic site, originally developed as anticancer drugs, such as silmitasertib (CX-4945) [19] [20, 21] and CIGB-300 [22] [23] have emerged as candidate for repurposing against COVID-19. A randomized and controlled clinical trial using CIGB-300 in COVID-19 patients with pneumonia demonstrated encouraging preliminary results [24]. Despite a small cohort of twenty patients, CIGB-300 treatment significantly reduced the median number of pulmonary lesions. However, ATP-competitive inhibitors suffer from a lack of selectivity, inhibiting several human kinases at nanomolar potency, which limits both their utility as a probe for studying CK2 function and their therapeutic potential.

Recently, AB668, a novel CK2 inhibitor, was introduced as a potent and highly selective molecule targeting CK2. Unlike traditional ATP-competitive inhibitors, AB668 binds simultaneously to the ATP site and the αD pocket of CK2α, thereby inhibiting the CK2 holoenzyme with high specificity [25]. Selectivity profiling against a panel of 468 kinases confirmed its minimal off-target activity, establishing AB668 as a valuable chemical probe to investigate CK2’s role in viral lifecycles. Moreover, its selectivity profile suggests a reduced likelihood of side effects caused by off-target kinase inhibition, enhancing its potential as a therapeutic agent.

In this study, we explore the implication of CK2α, CK2α’, and CK2β subunits in SARS-CoV-2 replication and their interaction with the SARS-CoV-2 nucleoprotein. We then evaluate the antiviral potential of AB668, a selective CK2 inhibitor, against SARS-CoV-2 and other coronaviruses, comparing its efficacy with that of CX4945. Finally, we assess its activity across multiple SARS-CoV-2 variants and its impact on different viral replication steps. These findings provide critical insights into the feasibility of targeting CK2 as a broad-spectrum antiviral strategy.

## MATERIALS AND METHODS

### Compounds

CX-4945, SGC-CK-2 and Remdesivir were purchased from Sigma-Aldrich. All compounds were dissolved in dimethyl sulfoxide (DMSO) at a final concentration of 0.1% in the experimental assays. AB668 was synthesized as previously published [26]. The HPLC-MS and NMR spectra are shown in supplementary information (Supplementary Figure 1).

### Viruses

SARS-CoV-2 ancestral strain B.117.18, Alpha and Delta variants were isolated from nasopharyngeal swabs collected at Henri Mondor University Hospital in Créteil, France, and propagated in Vero-TMPRSS2 cells. The Omicron B.A.1.529 variant was obtained from the European Virus Archive (EVAg, reference number 47184). HCoV-229E and HCoV-OC43 were sourced from Professor Astrid Vabret (CHU Caen, France) and the American Tissue and Culture Collection (ATCC, VR-1558), respectively, and propagated in MRC-5 cells (human lung fetal fibroblasts; ATCC CCL-171) and HRT-18 cells, respectively.

### Cell culture

Vero-TMPRSS2, Huh7, HEK-ACE2-TMPRSS2, and HRT-18 cells were cultured in Dulbecco’s Modified Eagle Medium (DMEM; Gibco™, Thermo Fisher Scientific®) supplemented with 10% fetal bovine serum (FBS), 50 IU/mL penicillin, 100 µg/mL streptomycin, and 0.1 µg/mL amphotericin B. For HRT-18 cells, the medium was further enriched with 10% sodium bicarbonate and 1% non-essential amino acids. HEK-ACE2-TMPRSS2 cells were maintained with additional supplements, including 100 µg/mL hygromycin, 1.5 µg/mL puromycin, and 2 µg/mL poly-D-lysine.

### Cellular viability

Vero-TMPRSS2 cells were seeded at 15,000 cells/well in 96-well plates and treated with increasing concentrations of CK2 inhibitors. After 24 hours, cell viability was quantified using the MTT assay. Briefly, 0.5 mg/mL of MTT (3-(4,5-dimethylthiazol-2-yl)-2,5-diphenyltetrazolium bromide was added to the culture medium, incubated, and the absorbance measured at 550 nm, with background absorbance subtracted.

### SiRNA

SiRNAs targeting the mRNA gene of CK2α (CNSK2A1), CK2α’ (CNSK2A2), and CK2β (CSNK2B) subunits were purchased from Thermo Fischer®. HEK-ACE2-TMPRSS2 cells (50,000/cells) were transfected with 15 nM siRNA using DharmaFECT^TM^ 4 reagent (Dharmacon^TM^). After 24 hours, cells were infected with SARS-CoV-2 ancestral B.117.18 strain at an MOI of 0.2. Supernatants were collected 24 hours post-infection for ELISA, western blot, TCID_50_, and viability assays, while cells were prepared for RT-qPCR, western blot, and immunofluorescence.

### Virus Inhibition Assay

Vero-TMPRSS2 cells (20,000 cells/well, 96-well plate), HEK-ACE2/TMPRSS2 cells (150,000 cells, 24-well plate), Huh7 cells (10,000 cells/well, 96-well plate) and HRT-18 cells (30,000 cells/well, 96-well plate) were infected with the indicated virus at an MOI of 0.2. After a 2-hour incubation at 37°C with 5% CO₂, cells were washed with PBS, and a fresh medium with CK2 inhibitors was added. Supernatants and cells were collected 24 hours post-infection for further analyses.

### TCID_50/ml_

Supernatants from siRNA-transfected and SARS-CoV-2-infected HEK-ACE2-TMPRSS2 cells were centrifuged (1,500 rpm, 5 minutes, 4°C) and used for serial dilutions on Vero-TMPRSS2 cells. Cytopathic effects were observed after 48 hours, and TCID_50_/ml values were calculated using the Reed-Muench method.

### Time-of-addition assay

Twenty thousand Vero-TMPRSS2 cells were infected with SARS-CoV-2 ancestral strain at an MOI of four. AB668 and CX-4945 compounds at a final concentration of 25 µM were added at different time points: 0h, 2h, 5h, 7h, 9h, and 12h post-infection. 24 hours post-infection, cells were washed with PBS and fixed for immunofluorescence assay or lysed for RT-qPCR measurement.

### Viral RNA Extraction and RT-qPCR

RNA from supernatants and cells were extracted using QIAamp Viral RNA Mini Kit (Qiagen) and RNeasy® Mini Kit (Qiagen), respectively. Reverse transcription was performed using the High-Capacity cDNA Reverse Transcription Kit (Thermo Fisher Scientific®). TaqMan Gene Expression Master Mix (Applied Biosystems) and GoTaq® Probe One-Step RT-qPCR System (Promega) were used for RT-qPCR. Relative RNA quantification employed the 2⁻ΔΔCT method, normalized to GAPDH or 18S rRNA.

### Western Blot

Protein lysates were prepared with EZ Lysis Buffer (20mM Tris HCL at pH 6.8, 100mM NaCl, 1mM EDTA, 0.5% NP40, and 10% Glycerol), and protein concentrations were determined using the Pierce BCA Protein Assay Kit (Thermo Fisher Scientific®). Samples were resolved on 4-12% Bis-Tris SDS-PAGE gels (BioRad) and transferred to nitrocellulose membranes. Membranes were incubated with primary antibodies against SARS-CoV-2 nucleoprotein (GeneTex, 1:1000) overnight at 4°C, followed by HRP-conjugated secondary antibodies. Signals were detected using ECL™ Prime Western Blotting Detection Reagents.

### Immunofluorescence

Vero-TMPRSS2 and HEK-ACE2-TMPRSS2 cells (500,000/well, 8-well Labtek Permanox plates) were infected with SARS-CoV-2 (MOI = 1) in the presence of CK2 inhibitors. Cells were fixed with 4% paraformaldehyde, permeabilized with 0.3% Triton X-114 and blocked with 10 mg/mL BSA. Immunostaining employed a primary antibody against SARS-CoV-2 nucleoprotein (GeneTex, 1:1000) and Alexa Fluor-conjugated secondary antibodies (Thermo Fisher Scientific, 1:1000).

### Co-Immunoprecipitation

CK2 subunit-specific antibodies (Santa Cruz), conjugated to magnetic beads, were used to immunoprecipitate CK2 subunits from lysates of HEK-ACE2-TMPRSS2 cells infected with SARS-CoV-2 (MOI = 1) according to manufacturer’s instructions (Dynabeads^TM^, Thermo Fisher scientific). The immunoprecipitates were then analyzed by western blot using an antibody directed against the SARS-CoV-2 nucleoprotein (Gentex). Uncoupled Magnetic beads were used as control.

### Bio-layer interferometry

CK2α (1-335 AA) was produced in Escherichia coli BL21 using plasmids kindly provided by Dr. Odile Filhol (Grenoble, France). CK2β (1-198) from plasmid pAB46 (addgen #27085) was subcloned into pGEX4-3T and expressed as GST-tagged protein in BL21 as previously described [26]. The interaction of the CK2α, CK2β and the holoenzyme CK2α2β2 with the SARS-CoV-2 nucleoprotein were analyzed via biolayer interferometry using the OctetRed96 instrument (ForteBio). Analyses were conducted at 30°C with constant shaking at 1,000 rpm in Kinetic buffer (KB: PBS, 0.02% Tween-20, 0.1% BSA). Briefly, his-tagged N protein of SARS-CoV-2 (R&D, 10474-CV; 10µg/mL in KB) was captured on anti-Penta-His (HIS1K) biosensors (Sartorius) during 600s. The biosensors loaded with SARS-CoV-2 N protein were dipped into wells containing 200nM CK2α, CK2β or CK2α2β2, and the association phase was monitored for 300s. Biosensors were then incubated further in KB for 300s to observe dissociation phase of the complex. Binding signals were obtained by subtracting the signals from empty HIS1K biosensor and from zero-concentration samples. For dissociation constant (KD) determination, SARS-CoV-2 N-loaded biosensors were dipped in wells containing 50, 100 or 200 nM CK2α2β2. Octet® Data Analysis software was used to determine KD values from the association and dissociation curves using a 1:1 binding model.

### Statistical Analysis

Data were analyzed using GraphPad Prism 9.0. Results are expressed as mean ± SD. Statistical significance was determined using Mann-Whitney U-tests or two-tailed Wilcoxon matched-pairs signed-rank tests. EC_50_ and CC_50_ values were derived using nonlinear regression. Significance levels were indicated as *p < 0.05, **p < 0.01, ***p < 0.001, or ns for non-significant differences.

## RESULTS

### Knockdown of CK2-Subunits (α, α’, and β)

The impact of siRNA targeting CK2α, CK2α’, and CK2β subunits on SARS-CoV-2 RNA production was assessed in HEK cells stably expressing ACE2 and TMPRSS2 (HEK-ACE2-TMPRSS2). The efficiency and specificity of siRNA-induced gene silencing was confirmed by RT-qPCR (Fig.S2). Twenty-four hours prior SARS-CoV-2 infection, cells were transfected with the indicated siRNA. Intracellular SARS-CoV-2 RNA level was quantified 24 hours post-infection using RT-qPCR. Knockdown of CK2α, CK2α’, and CK2β subunits significantly decreased SARS-CoV-2 intracellular RNA production by 64%, 61% and 74%, respectively (Fig.1A). Extracellular viral RNA production was similarly impaired (Fig.1A).

**Figure 1.**
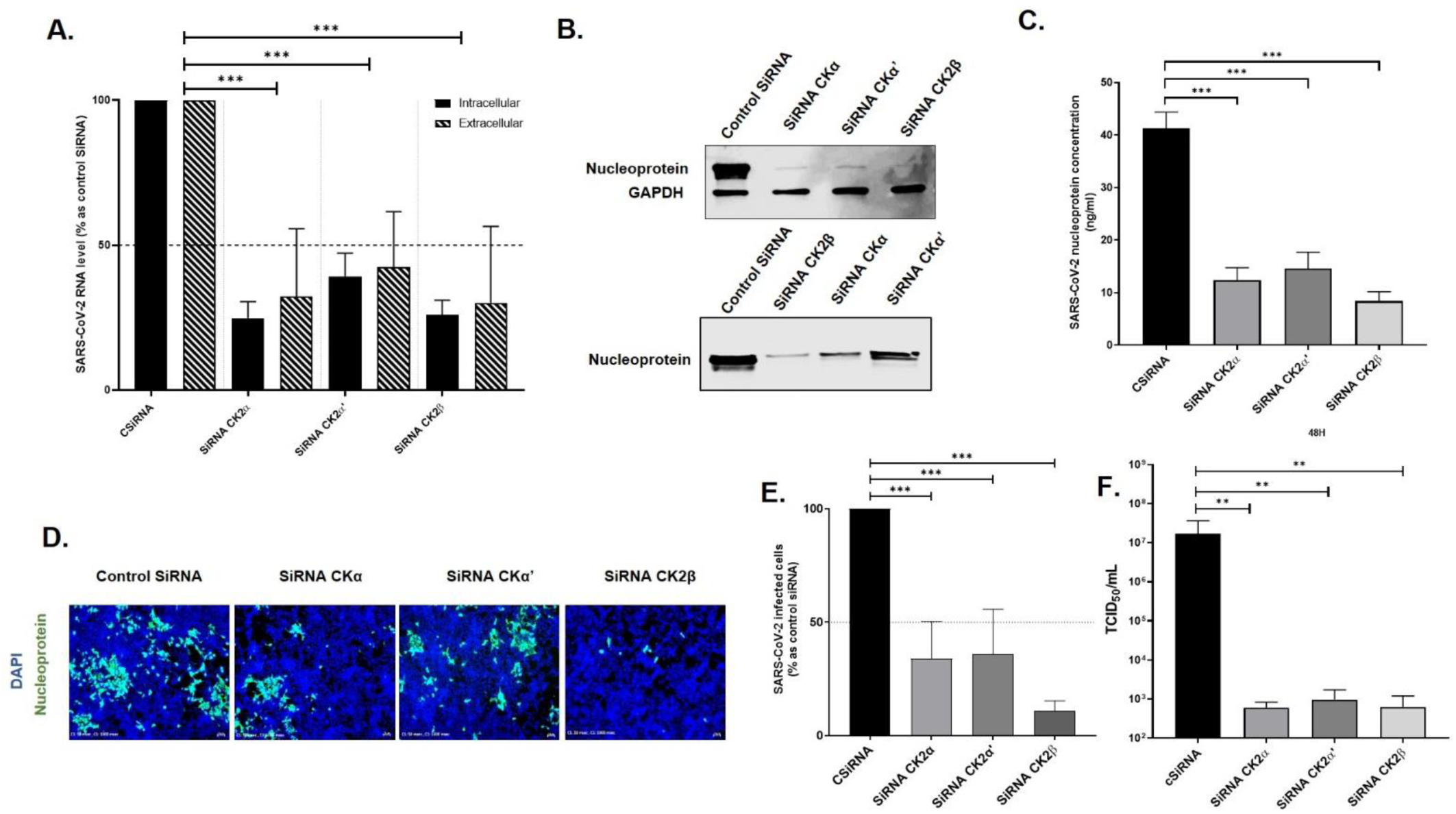
CK2 knockdown reduces SARS-CoV-2 replication in HEK-ACE2/TMPRSS2 cells. Following knockdown of CK2 subunits (α, α’, β) and SARS-CoV-2 infection, we performed**: (A)** Relative quantification of intracellular and extracellular SARS-CoV-2 RNA levels by RT-qPCR, expressed as % of Control SiRNA. **(B)** Analyze of intracellular and extracellular SARS-CoV-2 nucleoprotein levels by Western Blot; GAPDH was used as a loading control for protein normalization. **(C)** Quantification of nucleoprotein (ng/mL) release in infected cell supernatant by ELISA. **(D)** Immunofluorescence monitoring of SARS-CoV-2 infection (nucleoprotein in green, HEK-ACE2/TMPRSS2 nuclei in blue, 4× magnification) and **(E)** quantification of the number of infected cells (expressed as % of untreated control cells) using FIJI ImageJ software (22 fields per condition analyzed). **(F)** Determination of viral titer (log_10_ TCID_50_/mL). Statistical significance was determined using the Mann-Whitney test compared to the control siRNA values (*P<0.05, **P<0.01, ***P<0.001, ns = not significant).

The impact of siRNA targeting CK2α, CK2α’, and CK2β subunits was also assessed on SARS-CoV-2 nucleoprotein expression and release by means of western blot and ELISA. Similarly, knockdown of CK2α, CK2α’, and CK2β subunits highly reduced SARS-CoV-2 nucleoprotein expression (Fig1B, upper panel) and release in cell supernatant (Fig1B, lower panel) with a more pronounced effect for siRNA targeting CK2β and CK2α than CK2α’. Nucleoprotein release in cell supernatant was also quantified by ELISA with a 70%, 64% and 79% decreased of nucleoprotein concentration following CK2α, CK2α’, and CK2β siRNA treatment, respectively (Fig.1C).

The antiviral effect of siRNA targeting CK2α, CK2α’, and CK2β was further visualized through immunostaining of SARS-CoV-2 infected cells using an antibody targeting the nucleoprotein. A significant reduction in the number of infected cells was observed following knockdown of CK2α, CK2α’, and CK2β subunits (Fig.1D) with a more pronounced effect for CK2β. This decreased was quantified using image J software (Fig.1E).

Finally, the impact of CK2α, CK2α’, and CK2β silencing on viral titer was measured. As shown in figure 1F viral titer was seriously impaired resulting in approximately a 4-log reduction of TCID50/mL 48 hours post-infection.

### Pharmacological inhibition of CK2 kinase activity

The anti-SARS-CoV-2 impact of pharmacological inhibition of CK2 kinase activity was investigated in HEK-ACE2-TMPRSS2 cells and Vero-TMPRSS2 cells (Fig 2A and 2B, respectively). Two CK2 kinase inhibitors, CX4945 (ATP-competitive) and AB668 (bivalent), were tested. Both compounds demonstrated a dose-dependent reduction in intracellular SARS-CoV-2 RNA production, with a maximal decrease of 4-log and 2-Log for AB668 and CX4945, respectively. Importantly, neither AB668 nor Remdesivir exhibited cytotoxicity in our cellular models at the tested concentration (Fig. 2C). CX4945 exhibited 50 % cytotoxicity at concentration greater than20 µM. It is worth noting that the ATP-binding molecule SGC-CK-2, included as a control, was found to be cytotoxic at a concentration of 5 µM and was then excluded from subsequent analyses.

**Figure 2:**
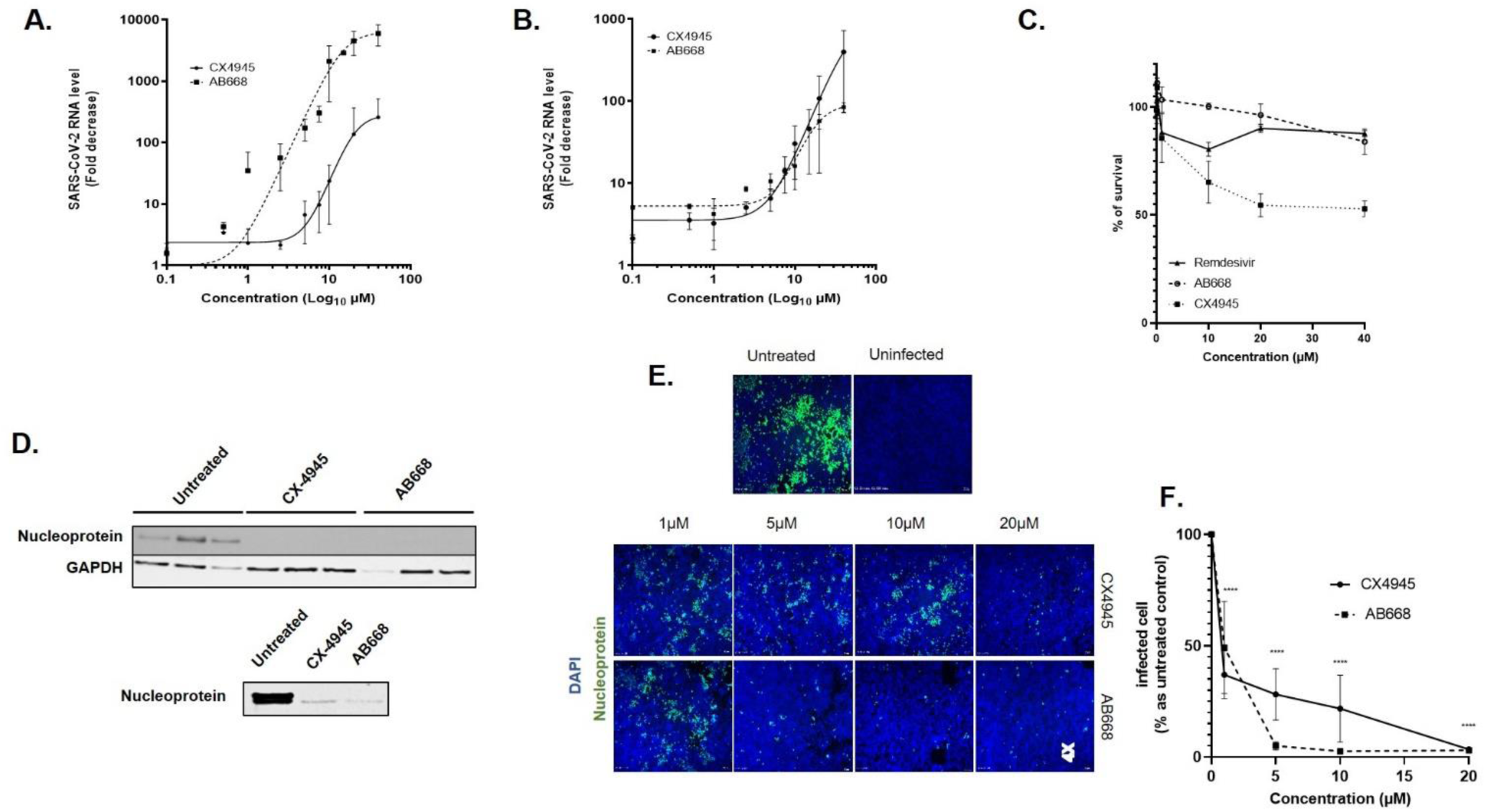
Pharmacological inhibition of CK2 kinase activity impaired SARS-CoV-2 replication. Relative SARS-CoV-2 RNA levels were quantified by RT-qPCR in **(A)** Vero-TMPRSS2 and **(B)** HEK-ACE2/TMPRSS2 cells treated with increasing concentration of CX-4945 and AB668. **(C)** Cell viability (expressed as % of untreated cell) was determined by MTT assay in HEK ACE2/TMPRSS2 cells. **(D)** Western Blot analysis of intracellular (upper panel) and extracellular (lower panel) of SARS-CoV-2 nucleoprotein in HEK-ACE/TMPRSS2 cells treated with 25 µM of CX4945 and AB668, GAPDH was used as a loading control for protein normalization. **(E)** Immunofluorescence staining of SARS-CoV-2 infected cells treated with the indicated concentration of CX4945 and AB668 (nucleoprotein in green, HEK-ACE2/TMPRSS2 nuclei in blue, 4× magnification **(F)** quantification of the number of infected cells (expressed as % of untreated control cells) using FIJI ImageJ software (12 fields per condition analyzed). The experiments were independently performed three times. Statistical significance was determined using the Mann-Whitney test compared to the control values. (*) represents P<0.05, (**) P<0.01, (***) P<0.001, (****) P<0.0001, and “ns” indicates no significance.

The impact of CK2 pharmacological inhibition on SARS-CoV-2 nucleoprotein expression and release was also assess by means of western blot analysis (Fig 2D). Both CX4945 and AB668 significantly reduced SARS-CoV-2 nucleoprotein expression (upper panel) and release (lower panel).

Furthermore, the antiviral effects of CX4945 and AB668 were evaluated using immunofluorescence. Staining of infected cells with an antibody targeting the SARS-CoV-2 nucleoprotein, revealed a dose-dependent decrease in the number of infected cells (Fig. 2E). The reduction of the number of infected cells was quantify using image J software (Fig. 2F) showing an EC_50_ of 0.8 ± 0.2 and 0.9 ± 0.3 µM for AB668 and CX4945, respectively

### CK2 holoenzyme interacts with SARS-CoV-2 nucleoprotein

The SARS-CoV-2 nucleoprotein is a well-characterized phosphoprotein, playing critical roles in the virus’s replication, transcription and assembly processes. Based on our previous findings that silencing of CK2 impairs SARS-CoV-2 replication, we sought to determine whether CK2 directly interacts with the nucleoprotein. To address this, HEK-ACE2-TMPRSS2 cells were infected with SARS-CoV-2 at a multiplicity of infection MOI of 1. Twenty-four hours post-infection, cell lysates were prepared, and CK2 subunits CK2α, CK2α’, and CK2β were immunoprecipitated using subunit-specific antibodies conjugated to magnetic beads. The immunoprecipitates were analyzed by western blot using an antibody against the SARS-CoV-2 nucleoprotein. As shown in Figure 3A, the nucleoprotein was strongly detected in the immunoprecipitates of CK2β while its presence was less pronounced in those of CK2α and CK2α’. These findings indicate an interaction between the nucleoprotein and CK2 subunits, particularly CK2β. To further characterize this interaction, we employed bio-layer interferometry (BLI) to measure the binding kinetics of the nucleoprotein with CK2α, CK2β, and the reconstituted holoenzyme CK2α2β2 *in vitro*. As shown in figure 3B, minimal or no interaction was observed between the nucleoprotein and the individual CK2α or CK2β subunits. However a stable and high-affinity interaction was detected between the nucleoprotein and the reconstituted holoenzyme CK2α2β2, with a dissociation constant (Kd) of 11.3 ± 0.3 nM (Fig. 3C). These results demonstrate that the nucleoprotein specifically interacts with the CK2 holoenzyme rather than its individual subunits, suggesting that the structural integrity of the holoenzyme is critical for this interaction

**Figure 3.**
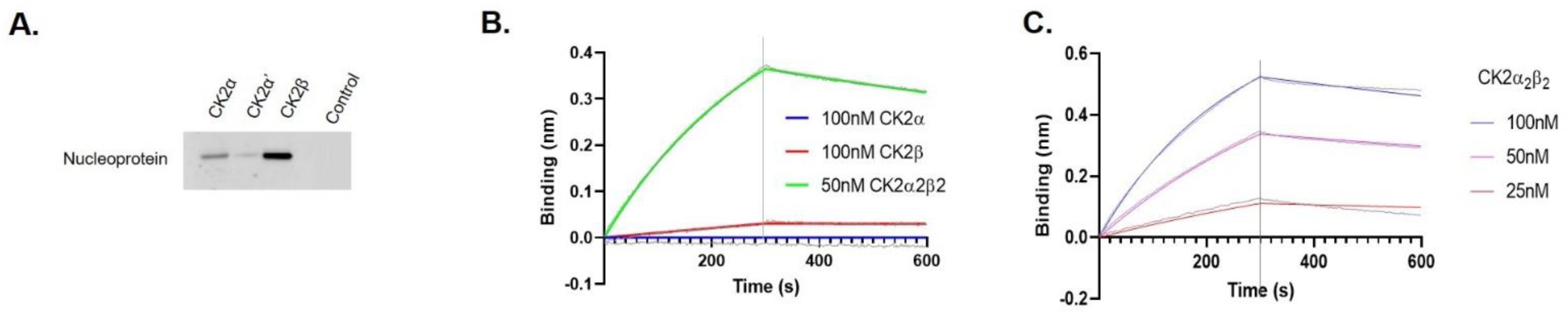
Interaction of CK2 with SARS-CoV-2 nucleoprotein. **(A)** Co-immunoprecipitation of SARS-CoV-2 nucleoprotein (NP) with CK2 subdomains (α, α’ and β) in HEK-ACE2/TMPRSS2 cells infected with SARS-CoV-2 (MOI = 1), visualized by Western Blot. **(B)** Biolayer interferometry (BLI) analysis of the interaction between NP-TagHis6 and CK2 subdomains or the holoenzyme. **(C)** Determination of the dissociation constant (Kd) values for the NP-CK2α’β2 interaction determined from BLI analysis. Experiments were independently repeated three times with consistent results.

### Viral step targeting by CK2 inhibitors

To identify the viral replication step targeted by CK2 inhibitors, a time of addition assay was performed using Vero-TMPRSS2 (Fig.4A). Cells were infected at a high multiplicity of infection (MOI = 4) to prevent multiple infection cycle. CK2 inhibitors (25 µM) were added at various time points relative to infection: at the time of infection (0 h) and at 2, 5, 7, 9, and 12 hours post-infection. At 24 hours post-infection, intracellular viral RNA and the number of infected cells were analyzed by immunofluorescence. The drug effect was calculated from fluorescence image using image J software (Fig. 4B). The results revealed that the drug effect decreased when added after 5 hours post infection with 50 % inhibitors lost antiviral activity when added 7 hours post-infection (Fig 4B). This suggests that the antiviral effects of these compounds target a step prior to or during SARS-CoV-2 RNA replication. To validate these findings, a single-cycle infection assay was performed in HEK-ACE2-TMPRSS2 cells using a lower MOI of 1 in the presence of CK2 inhibitors (25 µM). RT-qPCR analysis revealed a significant reduction in intracellular SARS-CoV-2 RNA levels 10 hours post-infection in the presence of both AB668 and CX4945 (Fig. 4C). These results confirm that AB668 and CX4945 exert their antiviral effects by targeting a step prior to or during the RNA replication phase in SARS-CoV-2 lifecycle.

**Figure 4.**
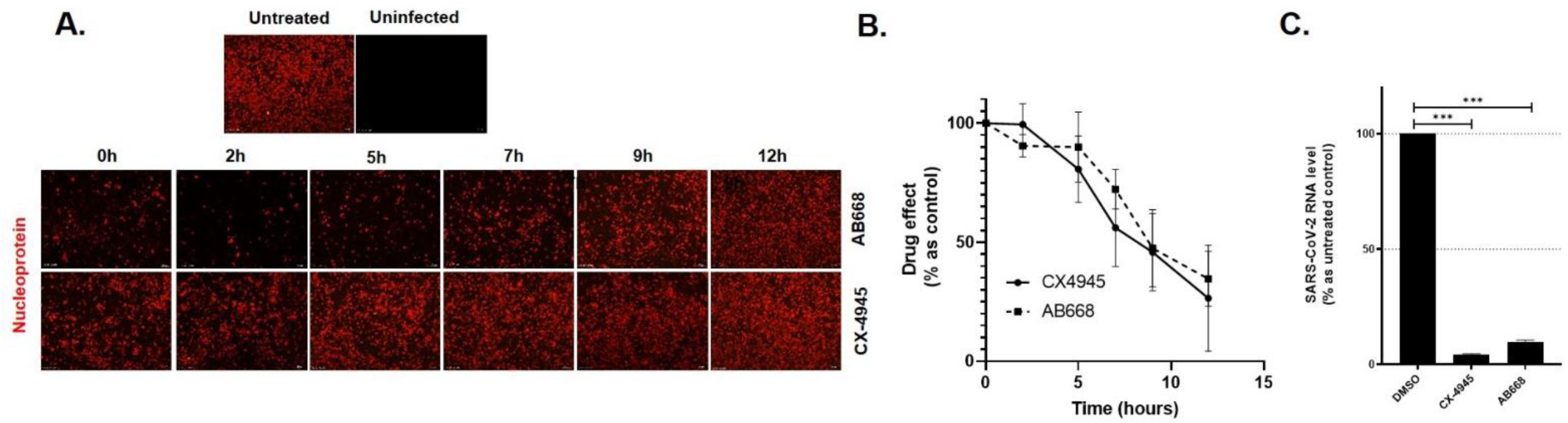
AB668 and CX-4945 inhibit SARS-CoV-2 at a pre or RNA replicative step. **(A)** Time-of-addition assay: Immunofluorescence staining of Vero-TMPRSS2 infected cells treated with AB668 or CX-4945 (25 µM) at 0, 2, 5, 7, 9 and 12 hours post infection. **(B)** The percentage of drugs effect was calculated from immunofluorescence image using FIJI ImageJ software. **(C)** Relative quantification of intracellular SARS-CoV-2 RNA production after 10-hours infection in HEK-ACE/TMPRSS2 cells treated with AB668 and CX-4945 (25 µM). The experiments were conducted independently three times in triplicates. All data represent the mean ± SD from three independent experiments performed in triplicate. Statistical analysis was performed using the Mann-Whitney test (*P<0.05, **P<0.01, ***P<0.001, ns = not significant).

### The Antiviral Spectrum of AB668 against Coronaviruses

The antiviral effectiveness of AB668 and CX-4945 against SARS-CoV-2 variants Alpha, Delta, and Omicron was assessed at concentrations of 10 µM and 20 µM in Vero-TMPRSS2 cells using RT-qPCR (Fig. 5A). The results indicated that both AB668 and CX-4945 exhibited lower efficacy against the Alpha variant compared to the ancestral strain. Neither compound showed significant antiviral activity against the Delta variant, while only AB668 demonstrated efficiency against the Omicron variant. These findings suggest differential CK2 dependency among SARS-CoV-2 variants. To further explore the role of CK2 in coronavirus replication, the antiviral effectiveness of AB668 and CX-4945 was tested against another betacoronavirus (OC43) and an alphacoronavirus (HCoV-229E) (Fig. 5B). The results showed that while both compounds were effective against the betacoronavirus OC43, only AB668 caused a dose-dependent reduction in RNA production of the alphacoronavirus HCoV-229E. These findings suggest that betacoronaviruses are more dependent on CK2 than alphacoronaviruses.

**Figure 5.**
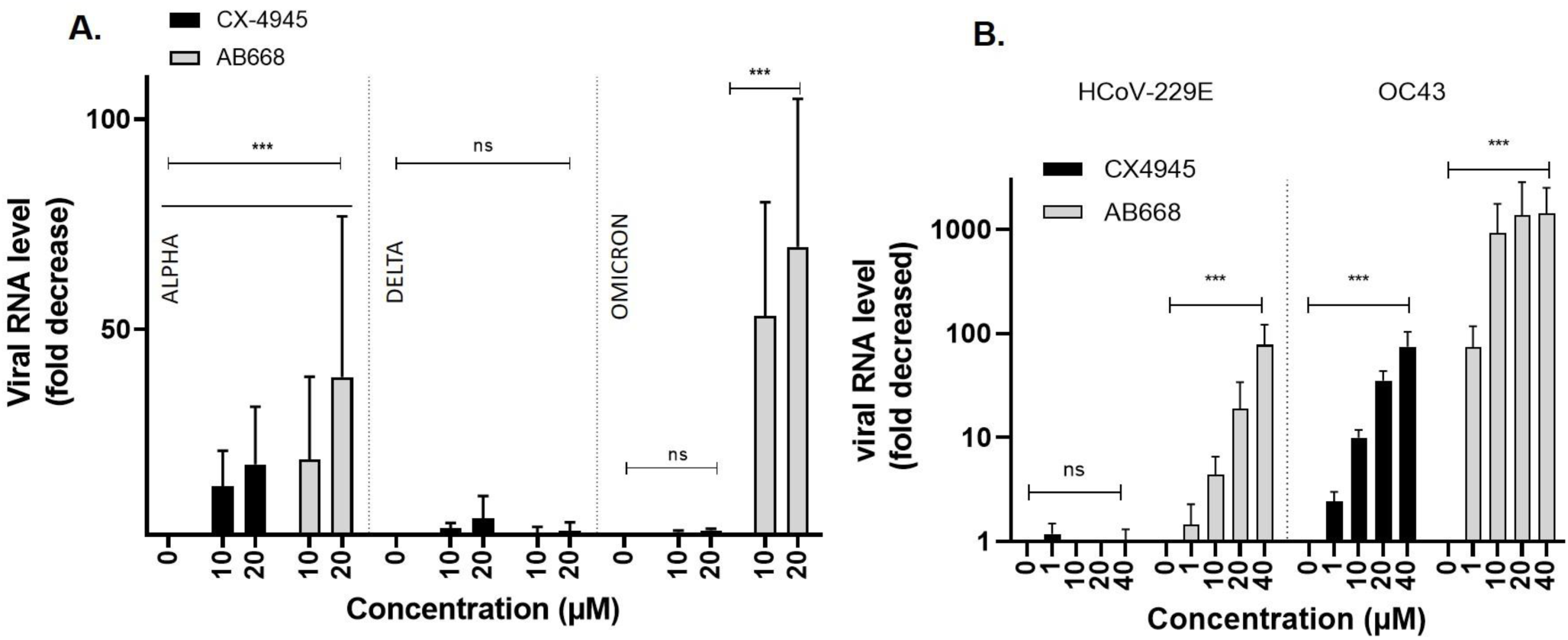
AB668 inhibits SARS-CoV-2 variants and other coronaviruses. **(A)** Fold decrease in intracellular RNA levels of SARS-CoV-2 variants measured by RT-qPCR in Vero-TMPRSS2 cells treated with 10 and 20 µM of AB668 or CX4945. **(B)** Antiviral activity of AB668 and CX-4945 against HCoV-229E and HCoV-OC43 evaluated by RT-qPCR. Intracellular viral RNA level is expressed as fold decrease compared to untreated controls. All experiments were performed in triplicates and independently repeated three times. Statistical analysis was carried out using the Mann-Whitney test (*P<0.05, **P<0.01, ***P<0.001, ****P<0.0001, ns = not significant).

In summary, these results demonstrate that AB668 possesses antiviral activity against multiple SARS-CoV-2 variants and exhibits broad-spectrum activity against coronaviruses, highlighting its potential as a pan-coronavirus antiviral agent.

## DISCUSSION

A comprehensive SARS-CoV-2 human interaction map identified 332 human protein interacting with 26 viral proteins. Among these, CK2 was mapped as interacting with the viral nucleoprotein [15]. Complementing this, a mass spectrometry-based phosphoproteomics study by Bouhaddou *et al*. revealed extensive rewiring of phosphorylation landscape in SARS-CoV-2-infected cells, and identified CK2 as a dysregulated host kinase [16]. Together, these global studies suggested a pivotal role for CK2 in SARS-CoV-2 lifecycle

Therefore, to confirm CK2 requirement to achieve SARS-CoV-2 replication and clarify the specific role of CK2α’, CK2α and CK2β subunits, targeted knockdown experiments were performed using subunits-specific siRNAs. The effectivity and specificity of the mRNA silencing were validated using RT-qPCR (Supplementary Fig2). Following siRNA-mediated knockdown, HEK-ACE2/TMPRSS2 cells were infected with SARS-CoV-2 and viral replication parameters, including viral RNA production, nucleoprotein expression, cellular infection and viral titer, were assessed. Knockdown of CK2α, CK2α’ and CK2β led to a significant decrease in all measured readouts, confirming that the CK2 tetrameric holoenzyme plays a critical role in SARS-CoV-2 replication. Interestingly, silencing of CK2α resulted in a slightly stronger effect compared to CK2α’, particularly with regard to nucleoprotein release in the cell supernatant. These findings suggest a preferential involvement of CK2α over CK2α’ in facilitating SARS-CoV-2 replication. To further investigate the interaction between CK2 subunits and the nucleoprotein in infected cells, co-immunoprecipitation (Co-IP) assays were performed. Magnetic beads were coated with specific antibodies against CK2α, CK2β, or CK2α’, as well as a combination of all three antibodies. The immunoprecipitated complexes were analyzed for nucleoprotein presence using Western blot. A significant co-imunoprecipitation of the nucleoprotein was observed when beads were coated with CK2α, CK2β, whereas a weak band was detected when beads were coated with CK2α’ antibody. These findings confirm the interaction between CK2 and the SARS-CoV-2 nucleoprotein and suggest a preferential binding of the nucleoprotein to CK2α compared to CK2α’. Finally, the interaction between the nucleoprotein and CK2α, CK2β, and the tetrameric CK2 holoenzyme (CK2α2β2) was evaluated using Bio-Layer Interferometry (BLI). While no significant interaction was observed between the nucleoprotein and CK2α or CK2β individually, a potent interaction was detected with the tetrameric CK2 holoenzyme reconstituted. This result demonstrates that the nucleoprotein interacts preferentially with the tetrameric CK2 holoenzyme rather than with the individual subunits alone. Overall, these findings indicate that the nucleoprotein interacts primarily with the tetrameric CK2 holoenzyme, with a preference for CK2α-containing holoenzymes.

Recently, AB668, a new generation of CK2 inhibitor has been developed. AB668 simultaneously bind both the ATP site and unique allosteric αD pocket on CK2, conferring outstanding selectivity over other kinases. Compared to CX-4945, the bivalent AB668 exhibited superior selectivity while minimizing off-target effects, addressing a major limitation of the ATP-competitive CK2 inhibitors.

The antiviral activity of AB668 against SARS-CoV-2 was extensively characterized. Our findings demonstrate that AB668 potently inhibits SARS-CoV-2 replication, significantly reducing viral RNA production, nucleoprotein expression and cellular infection across two different cell lines. Mechanistically, AB668 primarily acts at a post-entry step, within the first 7 hours post infection, suggesting its target is implicated in a pre-replicative or RNA replicative step. These results were further confirmed trough a single-cycle infection assay, which confirms that AB668 significantly reduced intracellular viral RNA production during the initial cycle of infection. Consistent with these findings, previous studies have suggested that the phosphorylation state of the nucleoprotein modulates critical processes, including protein-RNA interactions. Our results suggest that CK2 may play a critical role in regulating SARS-CoV-2 RNA replication by phosphorylating the nucleoprotein. Further studies are required to identify the specific residues on the nucleoprotein that are phosphorylated by CK2 and to elucidate the functional impact of these phosphorylation events on the viral lifecycle.

Expanding their SARS-CoV-2 human interaction map. Gordon *et al.* observed that CK2-nucleoprotein interaction was conserved across other β-CoV members, including SARS-CoV-1 and MERS-CoV [17], highlighting a potential role for CK2 in the β-CoV lifecycle. To validate this observation and to explore its relevance across other CoV genera, we assessed the antiviral effects of both CX-4945 and AB668 against SARS-CoV-2 variants Alpha, Delta and Omicron, as well as OC43 (β-CoV) and HCoV-229 (α-CoV).

Both CX4945 and the bivalent AB668 demonstrated inhibitory activity against the ancestral SARS-CoV-2 strain. However, their antiviral spectra varied among SARS-CoV-2 variants. AB668 showed efficacy against the Alpha and Omicron variants, while CX-4945 failed to inhibit Omicron variant replication. Neither compound demonstrated significant activity against the Delta variant. In agreement with the study of Bouhaddou et al. [27], our results suggest that SARS-CoV-2 variants evolve convergent strategies to remodel the host response, including altered nucleocapsid phosphorylation. Further studies are needed to map these interactions and analyze the phosphorylation profiles of the nucleoprotein in different SARS-CoV-2 variants to clarify these observations.

Regarding their activity against other human coronaviruses, both compounds effectively reduced HCoV-OC43 RNA production (a β-CoV). While AB668 was less efficient overall, it was still able, unlike CX4945, to significantly reduce HCoV-229E replication, an α-CoV. These results suggest that β-CoVs rely more heavily on CK2 activity than α-CoVs to achieve their replication. This differential dependence may be attributed to variations in the structural organization of their nucleoproteins, or the involvement of alternative host kinase pathways in α-CoVs. Detailed comparative studies on the role of CK2 across coronavirus families are needed to elucidate these differences. While both AB668 and CX4945 target CK2 their distinct binding modes and/or affinities likely contribute to the observed differences in antiviral activity. These findings underscore the need for further investigation into the mechanistic basis of their antiviral effects to optimize their therapeutic potential.

Overall, our findings establish AB668 as a promising candidate for broad-spectrum antiviral therapy. Its ability to selectively inhibit CK2 combined with efficacy against diverse SARS-CoV-2 variants and other β-CoV and α-CoV positions it as a valuable addition to the antiviral arsenal. Future studies should focus on evaluating AB668’s in vivo efficacy, pharmacokinetics, and potential for combination therapies.

## Supporting information

supplemental graph

supplemental LCMS

## REFERENCES

1. Venerando, A., M. Ruzzene, and L.A. Pinna, Casein kinase: the triple meaning of a misnomer. Biochem J, 2014. 460(2): p. 141–56.

2. Pinna, L.A., Protein kinase CK2. Int J Biochem Cell Biol, 1997. 29(4): p. 551–4.

3. Borgo, C., et al., Protein kinase CK2: a potential therapeutic target for diverse human diseases. Signal Transduct Target Ther, 2021. 6(1): p. 183.

4. Pinna, L.A., Protein kinase CK2: a challenge to canons. J Cell Sci, 2002. 115(Pt 20): p. 3873–8.

5. Montenarh, M. and C. Gotz, Protein Kinase CK2alpha’, More than a Backup of CK2alpha. Cells, 2023. 12(24).

6. Litchfield, D.W., et al., Functional specialization of CK2 isoforms and characterization of isoform-specific binding partners. Mol Cell Biochem, 2001. 227(1-2): p. 21–9.

7. Firzlaff, J.M., et al., The E7 protein of human papillomavirus type 16 is phosphorylated by casein kinase II. New Biol, 1989. 1(1): p. 44–53.

8. Meggio, F., et al., Phosphorylation of HIV-1 Rev protein: implication of protein kinase CK2 and pro-directed kinases. Biochem Biophys Res Commun, 1996. 226(2): p. 547–54.

9. Franck, N., et al., Hepatitis C virus NS2 protein is phosphorylated by the protein kinase CK2 and targeted for degradation to the proteasome. J Virol, 2005. 79(5): p. 2700–8.

10. Tang, J., et al., Phosphorylation of human La protein at Ser 366 by casein kinase II contributes to hepatitis B virus replication and expression in vitro. J Viral Hepat, 2013. 20(1): p. 24–33.

11. Davis, K.A., M. Morelli, and J.T. Patton, Rotavirus NSP1 Requires Casein Kinase II-Mediated Phosphorylation for Hijacking of Cullin-RING Ligases. mBio, 2017. 8(4).

12. Bajorek, M., et al., The Thr205 phosphorylation site within respiratory syncytial virus matrix (M) protein modulates M oligomerization and virus production. J Virol, 2014. 88(11): p. 6380–93.

13. Mitchell, C., et al., The nucleotidylylation of herpes simplex virus 1 regulatory protein alpha22 by human casein kinase II. J Biol Chem, 1997. 272(40): p. 25394–400.

14. Alvisi, G., et al., Multiple phosphorylation sites at the C-terminus regulate nuclear import of HCMV DNA polymerase processivity factor ppUL44. Virology, 2011. 417(2): p. 259–67.

15. Bouhaddou, M., et al., The Global Phosphorylation Landscape of SARS-CoV-2 Infection. Cell, 2020. 182(3): p. 685–712 e19.

16. Gordon, D.E., et al., A SARS-CoV-2 protein interaction map reveals targets for drug repurposing. Nature, 2020. 583(7816): p. 459–468.

17. Gordon, D.E., et al., Comparative host-coronavirus protein interaction networks reveal pan-viral disease mechanisms. Science, 2020. 370(6521).

18. Yang, X., et al., Host Kinase CSNK2 is a Target for Inhibition of Pathogenic SARS-like beta-Coronaviruses. ACS Chem Biol, 2022. 17(7): p. 1937–1950.

19. Pierre, F., et al., Pre-clinical characterization of CX-4945, a potent and selective small molecule inhibitor of CK2 for the treatment of cancer. Mol Cell Biochem, 2011. 356(1-2): p. 37–43.

20. Borgo, C. and M. Ruzzene, Protein kinase CK2 inhibition as a pharmacological strategy. Adv Protein Chem Struct Biol, 2021. 124: p. 23–46.

21. Chen, Y., et al., Strategies of Targeting CK2 in Drug Discovery: Challenges, Opportunities, and Emerging Prospects. J Med Chem, 2023. 66(4): p. 2257–2281.

22. Perea, S.E., et al., Antitumor effect of a novel proapoptotic peptide that impairs the phosphorylation by the protein kinase 2 (casein kinase 2). Cancer Res, 2004. 64(19): p. 7127–9.

23. Perera, Y., et al., *CIGB-300 anticancer peptide regulates the protein kinase CK2-dependent* phosphoproteome. Mol Cell Biochem, 2020. 470(1-2): p. 63–75.

24. Cruz, L.R., et al., Treatment with an Anti-CK2 Synthetic Peptide Improves Clinical Response in COVID-19 Patients with Pneumonia. A Randomized and Controlled Clinical Trial. ACS Pharmacol Transl Sci, 2021. 4(1): p. 206–212.

25. Bancet, A., et al., Cancer selective cell death induction by a bivalent CK2 inhibitor targeting the ATP site and the allosteric alphaD pocket. iScience, 2024. 27(2): p. 108903.

26. Grenier, D., et al., Binding-Site Switch for Protein Kinase CK2 Inhibitors. ChemMedChem, 2025: p. e202400868.

27. Bouhaddou, M., et al., SARS-CoV-2 variants evolve convergent strategies to remodel the host response. Cell, 2023. 186(21): p. 4597–4614 e26.

