## Supplementary figures and images for "Bivalent CK2 Inhibition as a Potent Strategy Against SARS-CoV-2 and Other Coronaviruses"

### supplemental graph

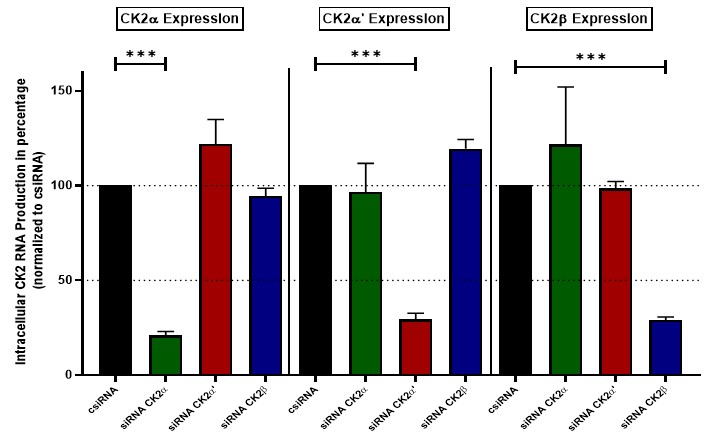

### supplemental LCMS

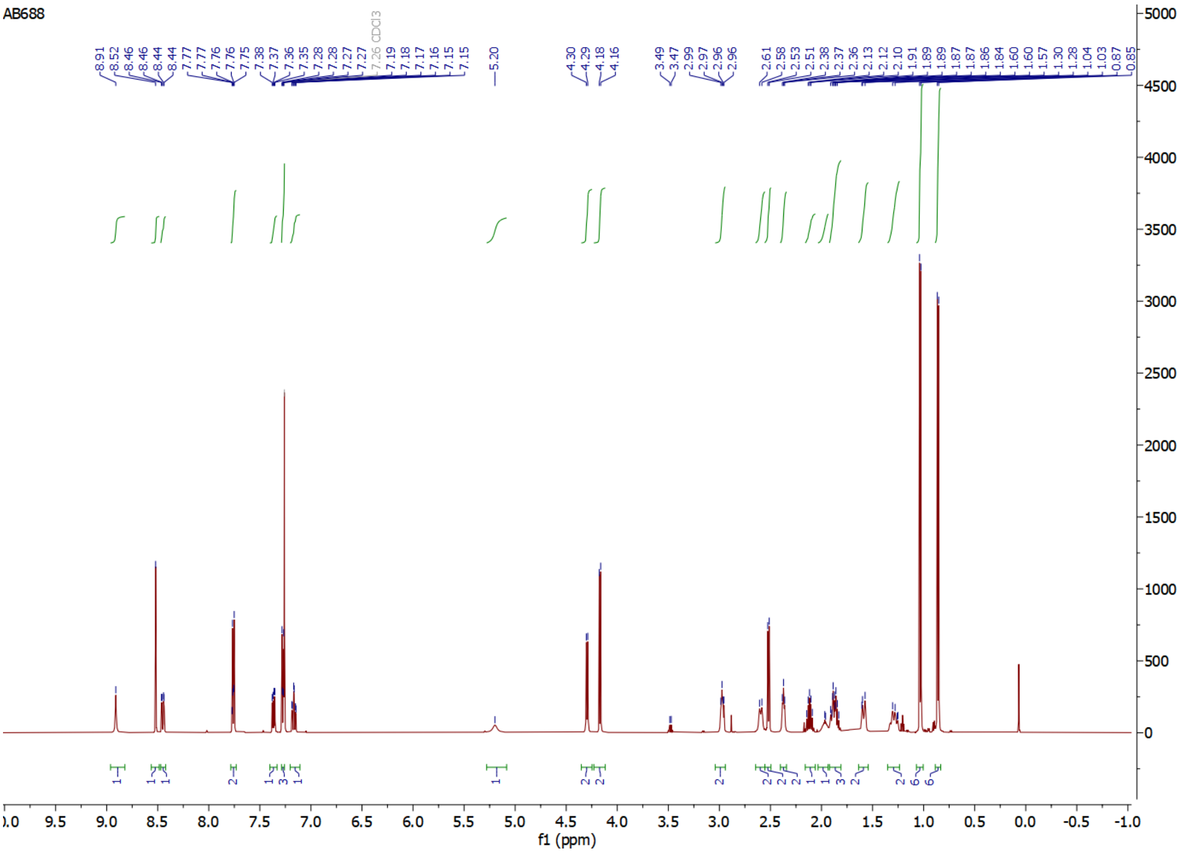

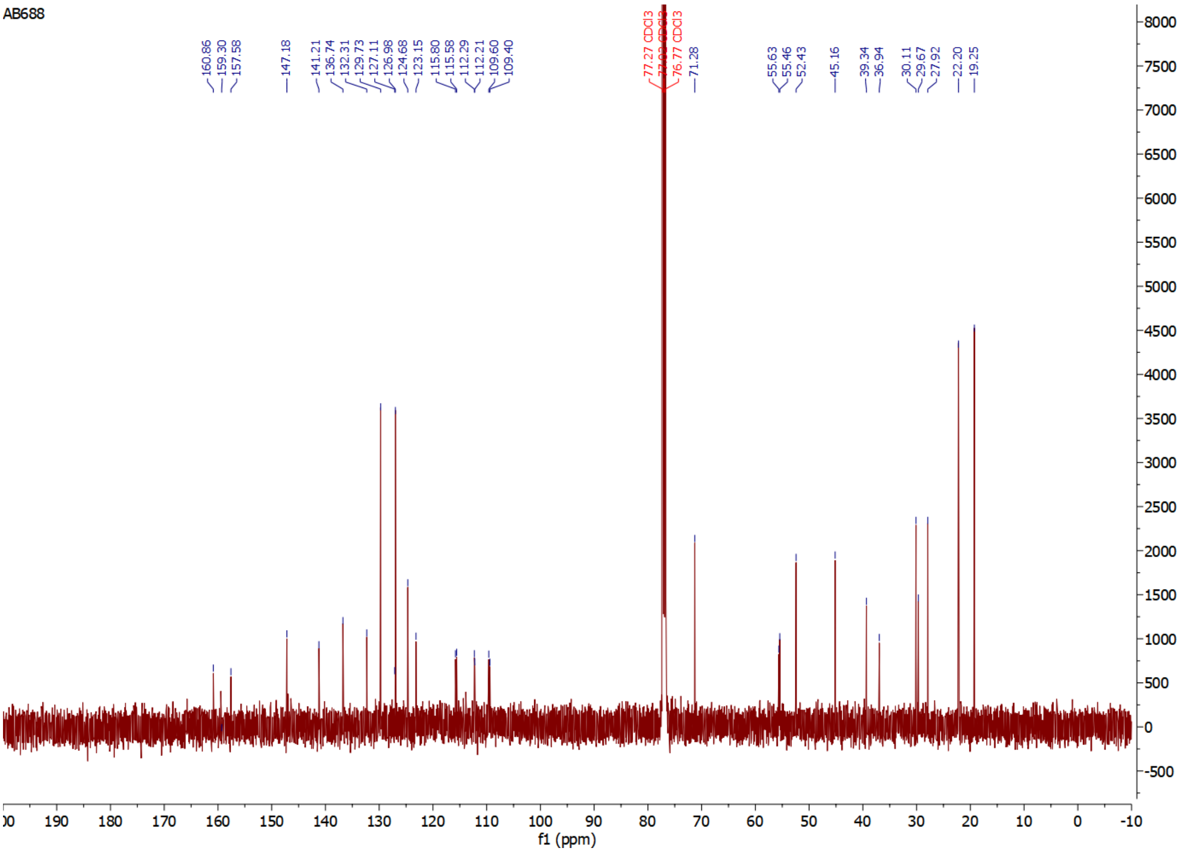

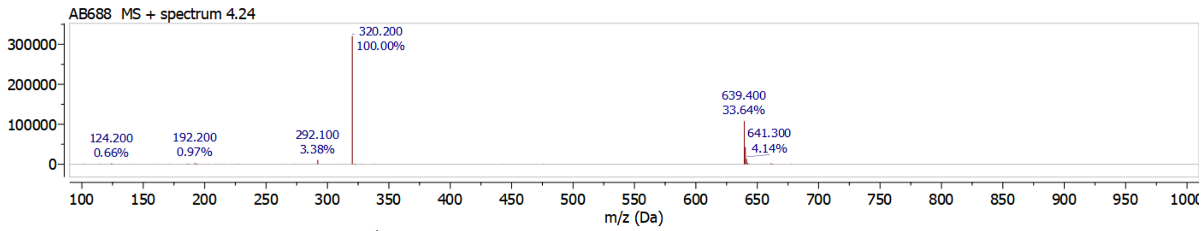


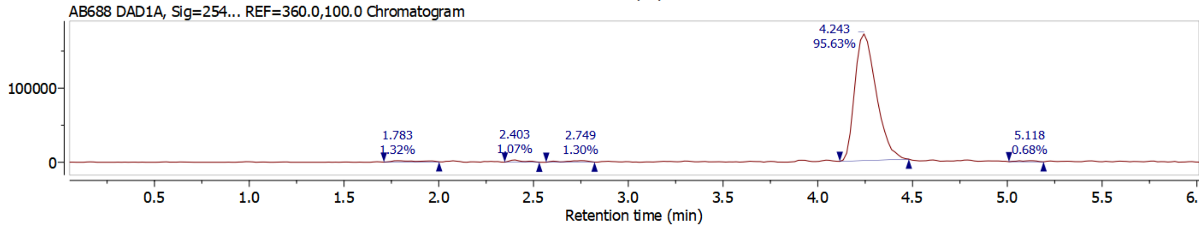


**Mass Peaks**

| | RT | Area | Total Height % | Total Area % |
|---|---|---|---|---|
| 1 | 1.783 | 17021.500 | 1.02 | 1.32 |
| 2 | 2.403 | 13755.000 | 1.66 | 1.07 |
| 3 | 2.749 | 16742.500 | 1.18 | 1.30 |
| 4 | 4.243 | 1231947.000 | 95.31 | 95.63 |
| 5 | 5.118 | 8822.500 | 0.83 | 0.68 |
